# Rod inputs arrive at horizontal cell somas in mouse retina solely via rod-cone coupling

**DOI:** 10.1101/2024.10.30.621116

**Authors:** Wallace B. Thoreson, Asia L. Sladek, Cody L. Barta, Lou E. Townsend

## Abstract

Rod and cone photoreceptor cells selectively contact different compartments of axon-bearing retinal horizontal cells in the retina. Cones synapse exclusively on the soma whereas rods synapse exclusively on a large axon terminal compartment. The possibility that rod signals can travel down the axon from terminal to soma has been proposed to allow spectrally opponent interactions between rods and cones, but there is conflicting data about whether this actually occurs. Because of spectral overlap between rod and cone visual pigments in mouse, we analyzed photoreceptor inputs into horizontal somata by selectively expressing channelrhodopsin in rods and/or cones. Optogenetic stimulation of rods and cones both evoked large fast inward currents in horizontal cell somata. Cone-driven responses were abolished by eliminating synaptic release in a cone-specific knockout of the exocytotic calcium sensor, synaptotagmin 1. However, rod-driven responses in horizontal somata were unchanged after eliminating synaptic release from rods but abolished by eliminating release from both rods and cones. This suggests that cones transmit rod signals that arrive via rod-cone gap junctions. Consistent with this, eliminating Cx36 gap junctions between rods and cones also abolished rod-driven optogenetic responses in horizontal cell somata. These results show that rod signals reach the somas of B-type horizontal cells exclusively via gap junctions with cones and not by traveling down the axon from the axon terminal.

**Significance:** Rods and cones contact different compartments of retinal horizontal cells: cones contact the soma whereas rods contact a large axon terminal. Cone signals can travel through the axon from soma to terminal, but it is not clear whether rod signals travel the other direction. This latter pathway has been proposed to mediate opponent interactions between rods and cones that could shape vision in dim lights. However, our results show that rod signals cannot travel from axon terminal to soma, but mix with cone signals via gap junctions between the two cell types. This finding limits mechanisms for explaining opponent rod-cone interactions in color and contrast perception.

## Introduction

Rod and cone photoreceptor cells transduce light into sensory receptor potentials and encode this information into patterns of synaptic vesicle release events (Moser et al., 2020; Thoreson, 2021). Synaptic release from photoreceptors is then shaped by lateral inhibition from second-order horizontal cells. Lateral inhibitory feedback from horizontal cells is essential for establishing center-surround receptive fields that enhance the detection of intensity gradients and provides a key substrate for color opponent interactions between different photoreceptors (Thoreson and Mangel, 2012). Rods and cones both synapse with horizontal cells and the possibility that rod-driven inputs into horizontal cells provide inhibitory feedback to cones has been invoked to explain several rod-cone interactions. This includes the presence of rod inputs in the inhibitory surround for JAMB OFF-center cells (Joesch and Meister, 2016), color opponency among RGCs in ventral regions of mouse retina (Szatko et al., 2020), and suppressive rod-cone interactions (Eysteinsson and Frumkes, 1989; Frumkes and Eysteinsson, 1988).

Most mammals have two types of horizontal cells: axonless A-type and axon-bearing B-type cells. In many species, axonless horizontal cells receive preferential or exclusive inputs from S cones. Mice have only axon-bearing horizontal cells (Peichl and Gonzalez-Soriano, 1994). Axon-bearing B-type horizontal cells have two anatomically distinct compartments: the soma and a large dendritic arbor at the axon terminal. Cones exclusively contact the soma and rods exclusively contact the axon terminal (Kolb, 1970; Kolb, 1974). Rod signals can enter cones via gap junctions but cone-driven light responses can be detected in axon terminals even in the absence of these gap junctions, showing they can also travel through the thin axon connecting the two compartments (Nelson et al., 1975; Steinberg, 1969; Trumpler et al., 2008). However, it is not clear if rod signals can travel via the axon in the other direction--from terminal to soma. This pathway is necessary for opponent interactions between rods and cones since rod signals that enter cones via gap junctions would mix with cone signals. Evidence that mouse cones may receive inhibitory feedback from rods thus supports the idea that rod signals can travel from the axon terminal compartment to the soma (Szikra et al., 2014). On the other hand, investigators failed to see rod-driven responses in horizontal cell somata from Cx36KO mice that lack gap junctions between rods and cones (Trumpler et al., 2008).

Given these conflicting results and the proposed role of this pathway in different mechanisms, we directly tested whether rod signals can travel down the axon from terminal to soma in mouse horizontal cells. To do so, we expressed channelrhodopsin2 (ChRh2) in rods or cones and recorded optogenetically-evoked responses in horizontal cell somata. We compared rod vs. cone-driven responses after genetically abolishing synaptic output from one or both photoreceptor cell types. We also tested optogenetic stimulation of rods in horizontal cells of mice that lack rod-cone gap junctions. Our results showed that while rod inputs can enter horizontal cell somata by traveling through gap junctions to cones, they are not transmitted through the axon from terminal to soma. This finding restricts mechanisms available for explaining opponent rod-cone interactions in the retina and visual system.

## Methods

### Mice

Mice were kept on 12 hour dark-light cycles. Mice aged 4-12 weeks of both sexes were used for experiments. Ai32 mice that express channelrhodopsin2/EYFP fusion protein in the presence of cre-recombinase were obtained from Jackson Labs. Rho-iCre (RRID:IMSR_JAX:015850) mice were also obtained from Jackson Labs (Li et al., 2005; Madisen et al., 2010). Details of HRGP-Cre and Syt1^flox^ (Syt1: MGI:99667) mice have been described previously (Le et al., 2004; Quadros et al., 2017). Syt1 was selectively eliminated from rods by crossing Syt1^fl/fl^ mice with Rho-iCre/Ai32 mice and eliminated from cones by crossing Syt1^fl/fl^ mice with HRGP-Cre/Ai32 mice. To eliminate Cx36, we originally tried breeding homozygous Cx36 global knockout mice (Deans et al., 2002) but learned after fruitless attempts that these mice are infertile. We next used Cx36^flox^ mice that allowed selective elimination of Cx36 gap junctions between rods and cones by crossing Cx36^fl/fl^ mice with Rho-iCre/Ai32 mice (Jin et al., 2022).

Euthanasia was conducted in accordance with AVMA Guidelines for the Euthanasia of Animals by CO_2_ asphyxiation followed by cervical dislocation. Animal care and handling protocols were approved by the institutional Animal Care and Use Committee.

### Electrophysiology

Mice were dark-adapted overnight and sacrificed in mid-morning. After euthanasia, retinas were isolated and horizontal slices of retina were prepared as described in detail elsewhere (Feigenspan and Babai, 2017). Briefly, retinas were i embedded in 1.8% low temperature gelling agarose (Sigma-Aldrich) and submerged in Ames’ medium supplemented with 5 mM HEPES. Horizontal slices (180-200 μm thick) were cut parallel to the plane of the retina using a vibratome (Leica Microsystems) at room temperature. Tissue slices were placed in the recording chamber and held in place with a slice anchor.

Tissue was superfused at 1-3 ml/min. with Ames supplemented with 5 mM HEPES and bubbled with 95% O2/ 5% CO2. The presence of HEPES limited effects of horizontal cell feedback during these recordings (Hirasawa and Kaneko, 2003). Whole cell recordings were obtained on an upright fixed-stage microscope (Olympus BX51 or Nikon E600FN) under a water-immersion objective (40x or 60x). We fabricated recording electrodes from borosilicate glass pipettes (1.2 mm OD, 0.95 mm ID, World Precision Instruments, Sarasota, FL) to produce electrodes with tip resistances of 10-12 MΩ. Electrodes were filled with potassium-based pipette solutions containing (in mM): 120 KSCN or KGluconate, 10 TEACl, 10 HEPES, 5-10 EGTA, 1 CaCl_2_, 1 MgCl_2_, 0.5 NaGTP, 5 MgATP, 5 phosphocreatine, 0.01 Alexa 488, pH 7.2-7.3. All chemical reagents were obtained from Sigma-Aldrich unless otherwise indicated.

Horizontal cells were identified visually and confirmed physiologically by their characteristic voltage-dependent currents, particularly prominent A-type K^+^ currents (Feigenspan and Babai, 2015). Horizontal cell identity was further confirmed in some cases by loading cells with the fluorescent dye Alexa 488 through the patch pipette.

Recordings were performed in voltage clamp using an Axopatch 200B (Axon Instruments/Molecular Devices) with AxoGraph X or PClamp10 acquisition software and digitized with an ITC-18 interface (Heka Instruments) or DigiData 1550 (Molecular Devices). Membrane currents were acquired at 10 kHz sampling and filtered at 1 kHz. Voltages were not corrected for liquid junction potentials (Gluconate pipette solution: 12 mV, KSCN pipette solution: 3.9 mV). Membrane capacitance, membrane resistance, and access resistance in horizontal cells averaged 16.9 ± 8.9 pF, 305.4 ± 252.1 MΩ, and 19.0 ± 4.7 MΩ (*n* = 41).

ChRh2 was activated by a 1-10 ms pulse of 490 nm light from an LED (Lambda TLED, Sutter Instruments). The voltage was regulated using a computer-controlled analog input to the LED. For experiments reported here, we used a voltage of 4V to generate consistently saturating responses.

Confocal images were obtained using Nikon Elements software and a laser confocal scanhead (Perkin Elmer Ultraview LCI) equipped with a cooled CCD camera (Hamamatsu Orca ER) mounted on a Nikon E600FN microscope. Fluorescent excitation was delivered from an argon/krypton laser at 488, 568, or 648 nm wavelengths and emission was collected at 525, 607, and 700 nm, respectively. Filters were controlled using a Sutter Lambda 10–2 filter wheel and controller. The objective (60X water immersion, 1.0 NA) was controlled using a E662 z-axis controller (Physik Instrumente). Images were examined and adjusted for color, brightness and contrast using Nikon Elements, Fiji/ImageJ, and Adobe Photoshop software.

### Statistical analysis

Statistical analysis and data visualization were done using GraphPad Prism. Where applicable, p values were adjusted for multiple comparisons using Tukey’s multiple comparisons tests along with oneway ANOVA. The criterion for statistical significance was set at α = 0.05. Data are presented as mean ± S.D

## Results

To compare rod and cone inputs into the soma compartment of mouse horizontal cell, we selectively expressed ChRh2 in rods and/or cones. To do so, we crossed Rho-iCre and HRGP-Cre mice that express Cre-recombinase in rods and cones, respectively, with Ai32 mice that express a Cre-driven channelrhodopsin-2 (ChRh2)/EYFP fusion protein. In HRGP-Cre/Ai32 and Rho-iCre/Ai32 mice that express ChRh2 in cones and rods, respectively, we used blue light to optogenetically stimulate photoreceptor cells. Using perforated patch recording techniques with gramicidin or beta-escin as a perforating agent, we found that optogenetic stimulation evoked depolarizing voltage changes in rods of 7.0 ± 3.54 mV (n=11) and cones of 6.6 ± 3.67 mV (n=4). Representative responses from a cone and rod are shown in Fig. 1A and B, respectively. Rod voltage response rose with a time constant of 8.8 ± 2.5 ms and reached their peak after 26.0 ± 13.6 ms. Cone responses were similar, attaining a peak response after 19.6 ± 8.8 ms.

**Fig. 1.**
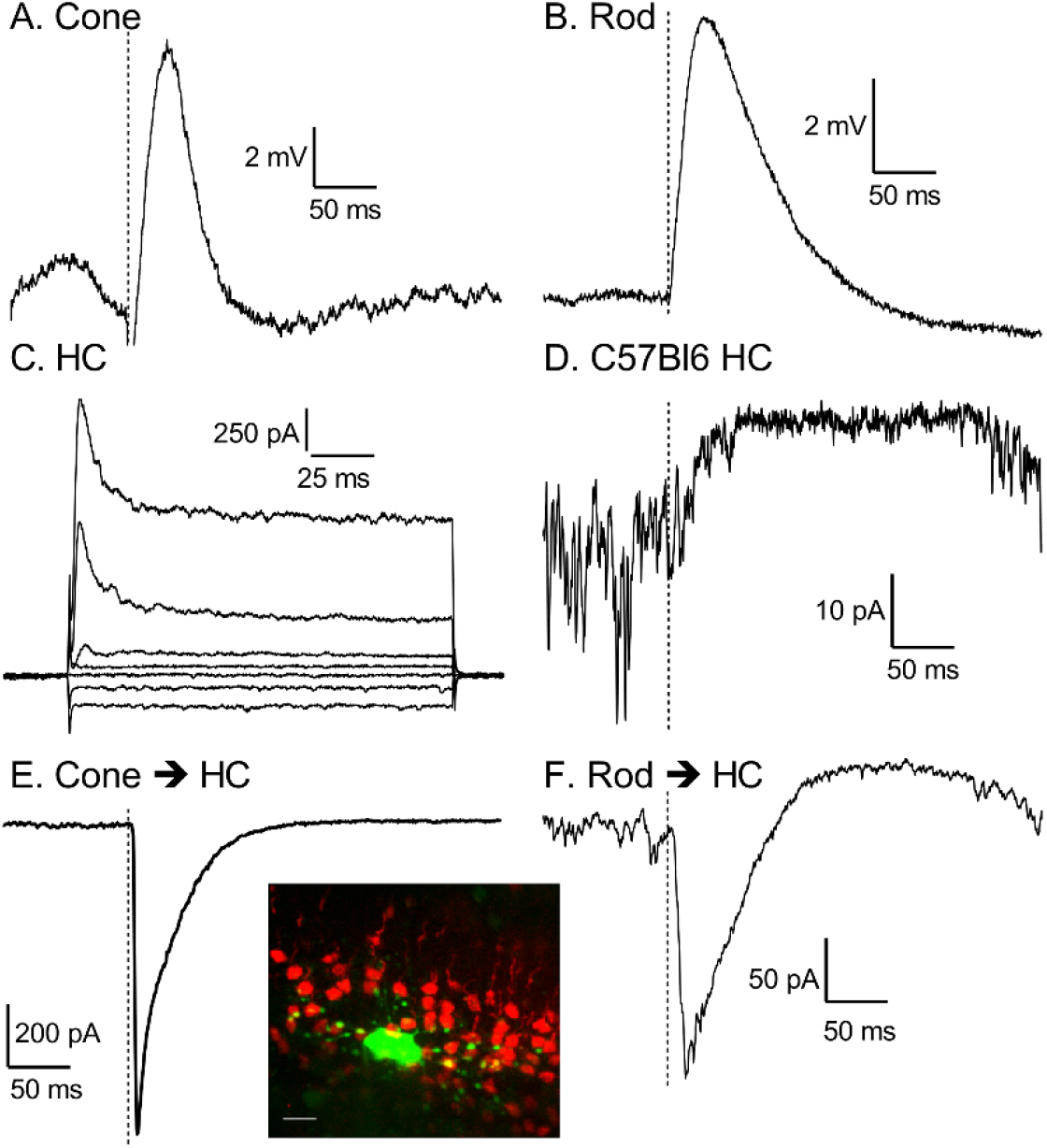
Responses of rods, cones, and horizontal cell somata to optogenetic stimulation of photoreceptor cells. A. Representative response of a cone expressing ChRh2 to 490 nm stimulation (dashed line). B. Response of a rod expressing ChRh2 to optogenetic stimulation (dashed line). C. Currents evoked by a series of voltage steps (20 mV steps, from -100 to +20 mV) applied to a horizontal cell. Prominent A-type outward currents are a characteristic of horizontal cells. D. Response recorded in a horizontal cell soma to a 1 ms 490 nm flash applied to a control B57Bl6J retina without ChRh2. E. Horizontal cell soma response to optogenetic activation of cones expressing ChRh2. Inset: horizontal cell from a control recording labeled with Alexa488 (green) along with cone terminals labeled by Cre-driven expression of Td-Tomato. F. Horizontal cell soma response to optogenetic activation of rods.

To record from horizontal cells, we prepared horizontal slices of retina, slicing along the plane of the inner nuclear layer to expose horizontal cells while retaining rod and cone inputs (Feigenspan and Babai, 2017). The fluorescence inset in Fig. 1 shows a horizontal cell filled with Alexa-488 in an HRGP-Cre retina where cones were labeled by Cre-driven expression of Td-tomato. Horizontal cells showed a characteristic set of voltage-dependent currents that included prominent A-type outward K^+^ currents (Fig. 1C; (Feigenspan and Babai, 2015)). In recordings from horizontal cells in retinas that lacked ChRh2, the bright blue light evoked slow outward currents (Fig. 1D) that arose from activation of endogenous cone opsins. By contrast with the slower kinetics of endogenous opsins, optogenetic stimulation of cones and rods that expressed ChRh2 evoked large, fast inward currents in horizontal cell somata (Fig. 1E and F). Endogenous opsins also contributed a late-developing outward current (e.g., Fig. 1F). Fig. 2 summarizes differences in amplitude and latency between rod- and cone-driven currents. The peak amplitude of the inward optogenetically-evoked current in horizontal cell somas was larger when driven by optogenetic stimulation of cones (565 ± 228 pA, n=27) than rods (291 ± 228 pA, p <0.0001, n=19). The latency to the peak of the inward current did not differ significantly between cone- (9.6 ± 2.80 ms; n=27) and rod- (11.2 ± 2.76 ms; n=19) driven responses. The peak of the horizontal cell current was achieved before the peak of the optogenetically-evoked voltage response in photoreceptors showing that synaptic release was efficiently triggered by the initial depolarization of these cells.

**Fig. 2.**
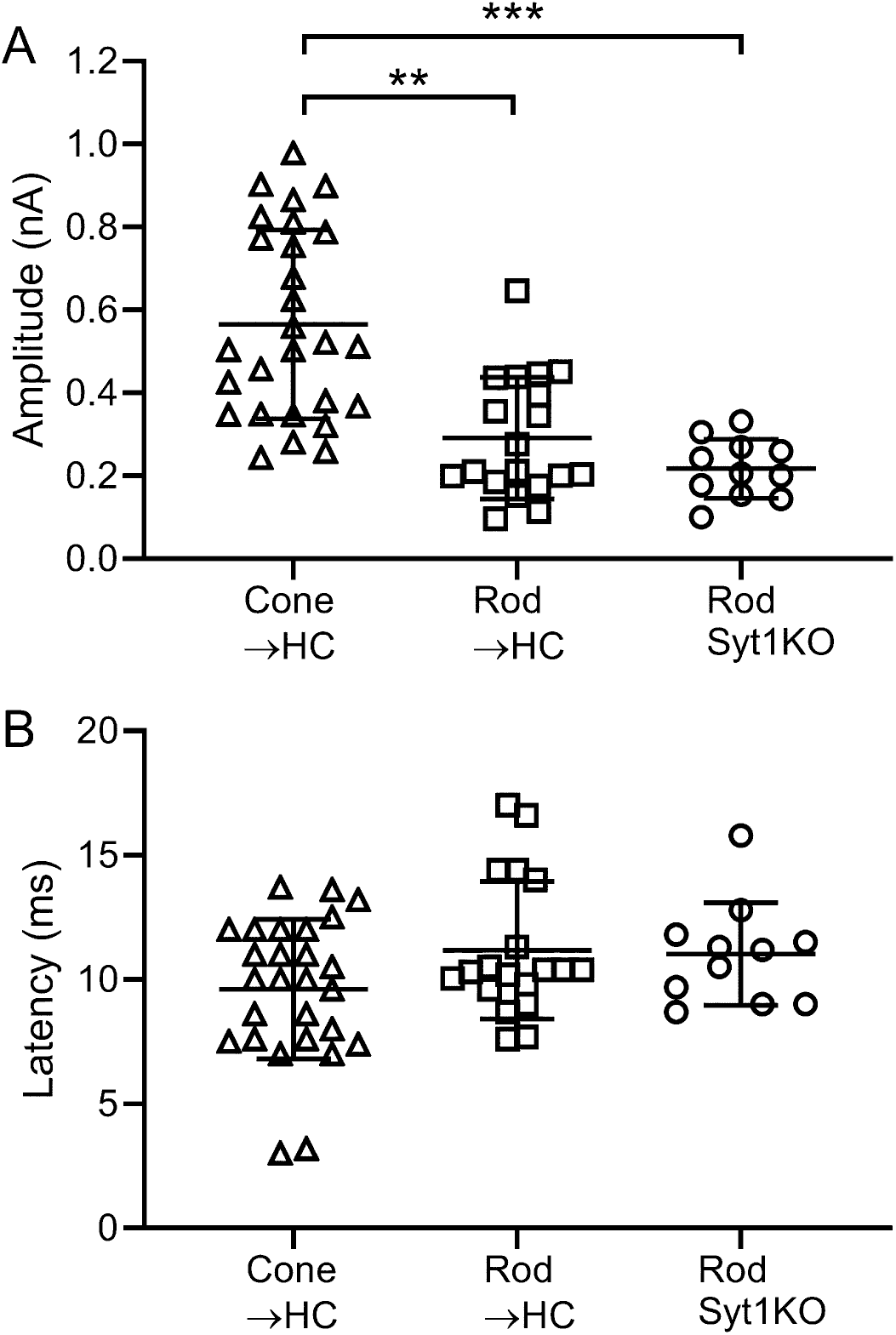
Summary data of current responses in horizontal cell somata evoked by optogenetic activation of cones (Cone → HC) and rods (Rod→HC). Data are also plotted for horizontal cell responses to optogenetic activation of rods that lacked Syt1 (Rho iCre/Syt1^fl/fl^). A. Peak amplitude of the inward optogenetically evoked current (± S.D.). Cone-driven: 565 ± 228 pA, n=27. Rod-driven: 291 ± 228 pA, n=19. Rod-driven/rod-specific Syt1KO: 217 ± 71.3 pA, n=11. ***, Cone-driven vs. rod-driven, p<0.0001, Cone-driven vs. rod-driven/rod-specific Syt1KO: p<0.0001, Tukey’s multiple comparisons test. B. Latency to the peak inward current did not differ among the three conditions. Cone-driven: 9.6 ± 2.80 ms. Rod-driven: 11.2 ± 2.76 ms. Rod-driven/rod-specific Syt1KO: 11.0 ± 2.06 ms.

To analyze pathways by which rod signals entered horizontal cell somata, we genetically eliminated the Ca^2+^ sensors controlling release in rods or cones. Fast glutamate release from both rods and cones is controlled by the presynaptic Ca^2+^ sensor, synaptotagmin 1 (Syt1) (Grassmeyer et al., 2019; Mesnard et al., 2022). As illustrated in Fig. 3A, selectively eliminating Syt1 from cones abolished horizontal cell responses to optogenetic stimulation of cones. Optogenetic responses were absent in all 7 horizontal cells recorded in 3 cone-specific Syt1 knockout retinas. These results support the conclusion that cone responses in the horizontal cell soma compartment arise from direct synaptic input from cones. Any cone responses that might enter rods via gap junctions do not appear capable of reaching the horizontal cell soma. While evoked responses were abolished, spontaneous miniature post-synaptic currents persisted in horizontal cells after loss of Syt1 in cones, consistent with results obtained previously by recording glutamate transporter currents in cones (Fig. 3A) (Grassmeyer et al., 2019).

**Fig. 3.**
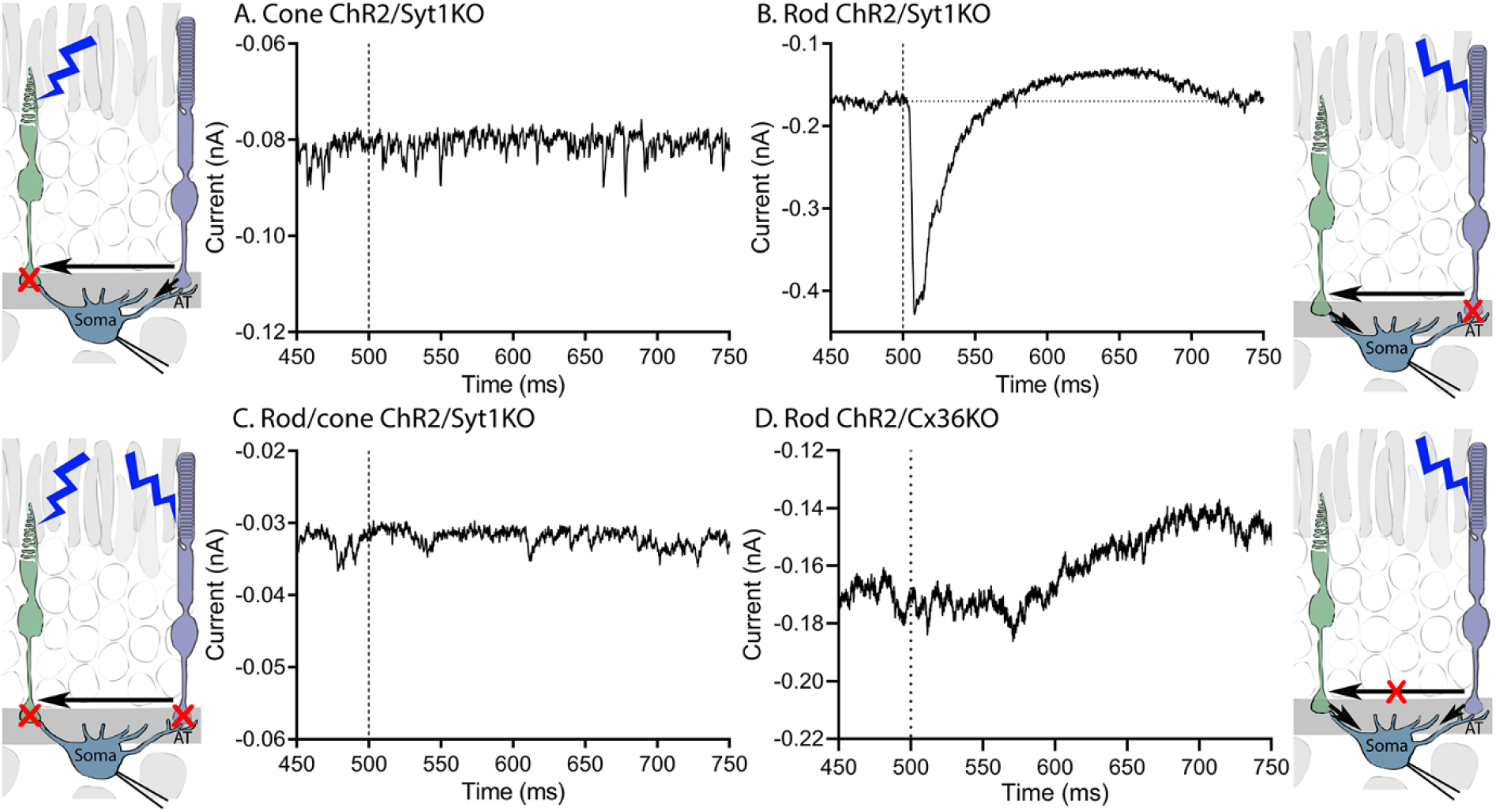
Rod signals enter horizontal somata via gap junctions with cones. A. Elimination of Syt1 from cones (Syt1^fl/fl^ x HRGP-Cre x Ai32) abolished their ability to evoke an optogenetic response (compare with Fig. 1E). B. Rods remained able to evoke large optogenetic responses in the absence of Syt1 (rod-specific Syt1KO: Syt1^fl/fl^ x Rho-iCre x Ai32). C. Eliminating Syt1 from both rods and cones (Syt1^fl/fl^ x HRGP-Cre x Rho-iCre x Ai32) eliminated responses to optogenetic stimulation of both photoreceptors in horizontal cell somata. D. Eliminating gap junctions with cones by removing Cx36 from rods (Cx36^fl/fl^ x Ai32 x Rho-iCre) abolished responses in horizontal cell somata evoked by optogenetic stimulation of rods.

By contrast with effects in cones, eliminating synaptic transmission from rods by genetically eliminating Syt1 did not significantly reduce currents evoked by optogenetic stimulation of rods (example in Fig. 3B, summary data in Fig. 2). The latency to peak current was also no different in mice with and without Syt1 in rods (Fig. 2).

We next eliminated Syt1 from both rods and cones. To do so, we crossed Syt1^fl/fl^ mice with Ai32 mice that express ChRh2 as well as HRGP-Cre and Rho-iCre mice that express cre-recombinase in cones and rods, respectively (Syt1^fl/fl^ x HRGP-Cre x Rho-iCre x Ai32). Since ChRh2 is expressed in both rods and cones, optogenetic stimulation activates both cell types. However, unlike the situation when synaptic transmission was only eliminated from rods, the additional loss of transmission from cones abolished horizontal cell responses to optogenetic stimulation (Fig. 3C; 5 cells from 3 mice). Once again, spontaneous release persisted in the absence of Syt1 (Fig. 3C). These data show that the entry of the rod-driven responses into horizontal cell somata does not involves synaptic release from rods and suggests that rod input instead reaches horizontal cell somata by transmission into cones via gap junctions.

To test the requirement for rod/cone gap junctions further, we recorded from mice that selectively lack gap junctions between rods and cones but retain Syt1 in rods (Cx36^fl/fl^ x Ai32 x Rho-iCre). In these mice, responses evoked by optogenetic stimulation of rods cannot be transmitted to cones via gap junctions and thus can only enter horizontal somata by transmission down the axon. Horizontal cells in these retinas exhibited mEPSCs and cone-driven light responses to the blue LED showing that they received functional synaptic input from cones. However, optogenetic stimulation of rods lacking gap junctions with cones abolished optogenetic currents in horizontal cell somata (Fig. 3D; n=8 cells from 3 mice).

## Discussion

The results of this study show that the entry of rod signals into horizontal cell somata requires both rod-cone gap junctions and intact synaptic output from cones. While rod signals can enter horizontal cell somata by traveling into cones via gap junctions from rods, they were abolished in the absence of rod-cone gap junctions and by eliminating Syt1 from cones. Our data thus support conclusions of Trumpler et al. (Trumpler et al., 2008) that cone signals can pass down the axon into the rod-driven horizontal cell terminal compartment, but rod signals cannot travel the other direction— from axon terminal to soma. The spectral sensitivity of horizontal cell somata therefore necessarily matches the spectral sensitivity of the cone input signal. Without a spectral difference, feedback from horizontal cell dendrites to cone somata cannot produce spectral opponency. Earlier recordings that showed evidence for rod to cone feedback were small and similar in size to extracellular field potentials that can be observed at that depth of the retina, raising the possibility that field potentials may have confounded interpretation (Szikra et al., 2014).

The asymmetry that allows cone signals to travel from soma to terminal but prevents rod signals from traveling the other direction involves asymmetries in input resistance, with a much higher input resistance in the axon terminal (Golard et al., 1992; Nelson et al., 1975; Yagi and Kaneko, 1988). Although post-synaptic currents will be attenuated by passage down the fine axon, small currents that reach a high resistance terminal compartment can generate measurable voltage changes. This is particularly evident in certain fish horizontal cells where the axon terminal compartment does not receive direct photoreceptor input but nevertheless shows cone-driven light responses (Stell, 1975; Weiler and Zettler, 1979).

Responses of rods and cones interact at many levels of the visual system. The earliest site involves gap junctions between rods and cones (Deans et al., 2002; Fain and Sampath, 2018; Jin et al., 2022; Sladek and Thoreson, 2023; Volgyi et al., 2004), allowing a pathway for rod signals to enter the cone circuitry. This so-called secondary pathway transmits rod signals more significantly at higher light intensities than the primary rod pathway involving direct contacts between rod and rod ON bipolar cells (Jin et al., 2022; Volgyi et al., 2004). Another early circuit by which rods and cones can interact involves inhibitory feedback from horizontal cells. Similar to cones, rods receive inhibitory feedback from post-synaptic horizontal cells (Babai and Thoreson, 2009; Thoreson et al., 2008). While our data argue against inhibitory feedback from rods to cones at horizontal cell somata, the presence of cone signals in the axon terminal compartment provides a substrate for inhibitory feedback of cone responses to rods. Mouse cones are more sensitive to short wavelengths than rods and so inhibitory feedback of mixed rod/cone signals in horizontal axon terminals to presynaptic rods would allow S/M wavelength opponent interactions.

The conclusions of this study limit potential sites available for opponent rod/cone interactions. For example, JAMB ganglion cells in mice show a center OFF response to UV light and surround ON response driven by rods (Joesch and Meister, 2016). Pharmacological experiments suggested that the surround response arose from horizontal cell feedback of rod signals to S cones. However, our data show that rod signals cannot travel into the soma via the axon, suggesting this pathway is unlikely to mediate the surround of JAMB cells and may instead involve inner retinal circuits. Similarly, horizontal cell feedback from rods to cones has been suggested as the mechanism mediating suppressive rod-cone interactions observed both psychophysically and at the cellular level in amphibians (Eysteinsson and Frumkes, 1989; Frumkes and Eysteinsson, 1988). In suppressive rod-cone interactions, the activity of rods in darkness suppresses cone-driven light responses. While feedback from rods to cones may provide such a mechanism in amphibians, our data argue that horizontal cell feedback from rods to cones is unlikely to be the mechanism for suppressive rod-cone interactions in mammals.

## Acknowledgements

We thank Dr. Christophe Ribelayga for generously providing Cx36^fl/fl^ mice used in these experiments. The authors declare no competing financial or other conflicts of interest. Funding provided by NIH grants EY10542 and EY32396 to WT.

